# AI-guided design of candidate BMPR1A-binding peptides for cartilage regeneration: a multi-tool computational benchmarking study

**DOI:** 10.64898/2026.03.22.713519

**Authors:** Asim Ahmadov, Olga Ahmadov

**Affiliations:** Department of Orthopaedics and Traumatology, Gazi University Faculty of Medicine, Ankara, Turkey; Department of Chest Diseases, Ankara University School of Medicine, Ankara, Turkey

## Abstract

Bone morphogenetic protein receptor type IA (BMPR1A) is a key mediator of chondrogenesis and a validated therapeutic target for cartilage repair, yet existing BMP mimetic peptides suffer from low potency and the full-length protein (rhBMP-2) carries significant safety risks. Generative AI tools for protein design can now produce de novo peptide binders, but none have been applied to cartilage regeneration targets. Here, we benchmarked four architecturally distinct AI tools—RFdiffusion, BindCraft, PepMLM, and RFpeptides—to design candidate BMPR1A-binding peptides. We generated 192 candidates alongside 98 negative controls (290 total) and evaluated all complexes using AlphaFold 3 structure prediction, dual physics-based energy scoring (PyRosetta and FoldX), and contact recapitulation against the crystallographic BMP-2:BMPR1A interface (PDB: 1REW). A four-metric composite ranking identified a 15-residue PepMLM design (pepmlm_L15_0026) as the top candidate, combining favorable binding energy (PyRosetta *dG*_separated_ = −45.9 REU; FoldX Δ*G* = −19.4 kcal/mol) with the highest contact recapitulation among top-ranked peptides (11/30 gold-standard interface residues). Designed candidates significantly outperformed controls on ipTM (*p* = 0.002) and FoldX Δ*G* (*p* < 0.001). BindCraft candidates achieved the highest structural confidence (ipTM up to 0.81) but exhibited moderate contact recapitulation (mean 0.224), consistent with the computational hypothesis that they may engage alternative BMPR1A binding surfaces rather than the native BMP-2 interface. Physicochemical filtering yielded a shortlist of 54 candidates across all four tools. These results establish a reproducible computational framework for AI-guided peptide design targeting cartilage regeneration and identify specific candidates for future experimental validation via binding assays and chondrocyte differentiation studies.

**Author summary:** Damaged cartilage has limited capacity to heal, and current biological therapies based on bone morphogenetic protein 2 (BMP-2) carry serious safety concerns including ectopic bone formation and inflammation. Short peptides that mimic BMP-2’s interaction with its receptor BMPR1A could offer a safer, more targeted alternative, but designing such peptides from scratch is challenging. We used four different artificial intelligence tools—each employing a distinct computational strategy—to generate 192 candidate peptides designed to bind BMPR1A. We then evaluated all candidates using multiple independent computational methods to assess binding quality, energy favorability, and whether each peptide targets the correct site on the receptor. Our analysis identified a shortlist of 54 promising candidates, with a 15-residue peptide from the language model-based tool PepMLM emerging as the top-ranked design. We also found evidence that one tool (BindCraft) may produce peptides that bind BMPR1A at sites different from the natural BMP-2 interface, highlighting the importance of validating not just whether a peptide binds, but where it binds. Our computational framework and candidate peptides provide a foundation for future laboratory testing toward cartilage repair therapies.

## Introduction

Articular cartilage has limited intrinsic repair capacity: lesions generally do not heal spontaneously, and effective natural repair occurs only within a narrow defect size range, owing in part to the tissue’s avascular nature [1]. Bone morphogenetic protein 2 (BMP-2), first cloned by Wozney et al. [2], is a potent inducer of both bone and cartilage formation in vivo. BMP-2 signals through heterotetrameric receptor complexes comprising type I and type II serine/threonine kinase receptors, activating the canonical Smad1/5/8 pathway in chondrocytes [3]. Among the type I receptors, BMPR1A (ALK3) has emerged as a critical mediator of chondrogenesis: conditional knockout studies demonstrate that BMPR1A is necessary for both chondrogenic and osteogenic differentiation [4], while the closely related BMPR1B acts as a hypertrophy suppressor [5, 6]. ACVR1 (ALK2) coordinates with BMPR1A and BMPR1B to regulate endochondral ossification [7]. Clinically, BMPR1A expression in human articular cartilage correlates with osteoarthritis (OA) progression, declining as cartilage degenerates [8]. The therapeutic potential of targeting this receptor was demonstrated by Akkiraju et al., who showed that CK2.1—a designed BMPR1A mimetic peptide—achieved complete repair of articular cartilage in an OA mouse model without inducing chondrocyte hypertrophy [9].

Despite this biological rationale, clinical translation of BMP-2 itself has been problematic. Recombinant human BMP-2 (rhBMP-2, marketed as INFUSE) was approved for spinal fusion, but a critical review of industry-sponsored trials revealed that adverse event rates were 10–50-fold higher than originally reported, including ectopic bone formation, radiculitis, osteolysis, and an apparent dose-dependent increase in malignancy risk [10]. Subsequent studies documented neurological impairment and ectopic ossification in cervical spine applications [11, 12], dose-dependent complication rates from FDA adverse event databases [13], and a broad spectrum of surgical complications across indications [14]. These safety failures underscore the need for alternatives that activate BMP signaling at lower effective doses with greater target specificity.

BMP mimetic peptides represent one such alternative. Short peptides derived from the BMP-2 “knuckle” epitope—the receptor-binding loop that contacts BMPR1A—have been shown to promote bone formation when delivered from controlled-release matrices [15, 16]. More recent work has incorporated BMP-2 mimetic peptides into hydrogel and cryogel scaffolds for cartilage and bone applications [17, 18]. However, these naturally derived peptides suffer from a fundamental limitation: even at 12,000-fold higher molar concentrations, BMP-2 knuckle peptides exhibit significantly lower osteoinductive potential than the full-length protein, partly due to peptide aggregation and suboptimal receptor engagement [19]. Designing peptides that bind BMPR1A with higher affinity than natural BMP-2 fragments requires exploring sequence space beyond what evolution has sampled.

Recent advances in generative AI for protein design offer precisely this capability. RFdiffusion generates protein backbones through denoising diffusion on a structure prediction network and has been extended to design high-affinity peptide binders [20, 21]. BindCraft performs one-shot binder design guided by AlphaFold 2 multimer confidence [22]. PepMLM conditions a protein language model on the target sequence to generate binding peptides without requiring structural information [23]. RFpeptides applies denoising diffusion specifically to macrocyclic peptide binder design [24]. Together, these tools represent four distinct architectural paradigms—diffusion-based, structure-guided, language model, and macrocyclic—each with potentially complementary strengths [25, 26]. Despite their demonstrated success on benchmark targets, no study has applied these generative AI tools to design peptides for cartilage regeneration.

Here, we present the first computational benchmarking study of AI-designed peptides targeting the BMPR1A extracellular domain for cartilage repair. We used all four tools to generate 192 candidate peptides, validated them alongside 98 negative controls (290 total) using AlphaFold 3 structure prediction [27], dual physics-based energy scoring (PyRosetta and FoldX), and contact recapitulation against the gold-standard BMP-2:BMPR1A crystal structure (PDB: 1REW) [28]. Our four-metric composite ranking identifies top candidates for future experimental validation and reveals distinct design trade-offs across AI architectures.

## Materials and methods

### Computational environment

Local analyses were performed in Python 3.11 using BioPython (v1.86), SciPy (v1.13), NumPy (v1.26), pandas (v2.1), matplotlib (v3.8), and seaborn (v0.13). AI peptide generation tools (PepMLM, RFdiffusion, BindCraft, RFpeptides) were executed on Google Colab (Python 3.10, NVIDIA T4 or L4 GPUs) using the default package versions provided by each tool’s official notebook at the time of execution (March 2026). FoldX v5.1 (build 20270131) was run locally on macOS ARM.

### Target preparation

The crystal structure of the BMP-2:BMPR1A extracellular domain (ECD) binary complex (PDB: 1REW) [28] was retrieved from the RCSB Protein Data Bank [29] as the primary target structure. Three additional structures were obtained for context and specificity analysis: the BMP-2:BMPR1A:ActRII ternary complex (PDB: 2H62), the BMPR1B ECD with BMP-2 (PDB: 3EVS), and an ACVR1 structure (PDB: 3MTF). Individual chains were extracted using BioPython: the BMPR1A ECD (86 residues) from chain C of 1REW served as the target for all peptide design tools.

A gold-standard interface was defined by computing inter-chain atomic contacts at a 5.0 Å cutoff between BMP-2 (chain A) and BMPR1A (chain C) in 1REW using BioPython’s NeighborSearch algorithm. This identified 30 BMPR1A interface residues and 20 BMP-2 interface residues. These residues defined the hotspot configurations exported in tool-specific formats for each AI design method.

### AI peptide generation

Four generative AI tools representing distinct architectural paradigms were used to design candidate BMPR1A-binding peptides (Table 1):

**Table 1.**
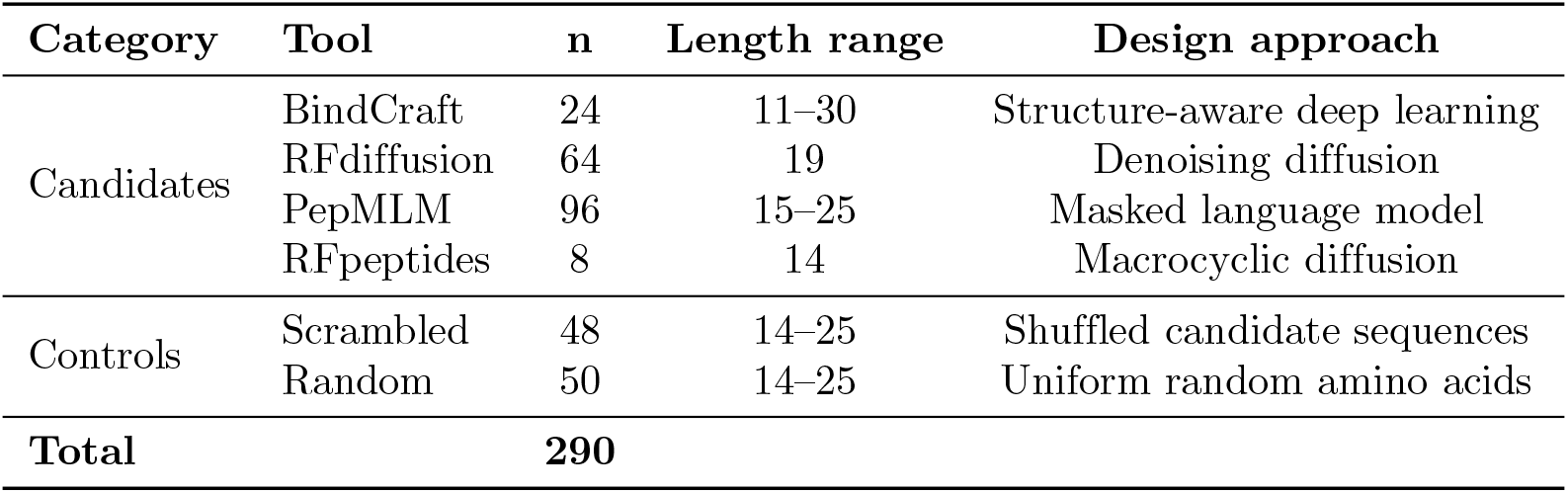
Study design overview. Four generative AI tools representing distinct architectural paradigms were used to design BMPR1A-binding peptide candidates. Controls include sequence-scrambled versions of designed peptides (preserving amino acid composition) and fully random sequences (uniform amino acid sampling at matched lengths).

**PepMLM** [23] (model: ChatterjeeLab/PepMLM-650M) is a target sequence-conditioned peptide generator that fine-tunes the ESM-2 protein language model using a span masking strategy. It requires only the target protein sequence as input, with no structural information. We generated 96 candidates at three peptide lengths (32 each at 15, 20, and 25 residues) conditioned on the BMPR1A ECD sequence.

**RFdiffusion** [20, 21] generates protein backbones through denoising diffusion on the RoseTTAFold structure prediction network. Peptide backbones were generated with hotspot residue guidance toward the BMPR1A interface. Amino acid sequences were then designed onto each backbone using ProteinMPNN [30] with default sampling temperature and no additional backbone noise, yielding 64 candidates (8 backbones × 8 sequences per backbone) of 19 residues each.

**BindCraft** [22] performs one-shot protein binder design guided by AlphaFold 2 [31] multimer predictions. The peptide-optimized three-stage multimer protocol was used with peptide-specific filters, targeting 10–30 residue binders against the BMPR1A hotspot residues. From 315 design trajectories run on Google Colab Pro (L4 GPU), 24 candidates (7.6% acceptance rate) passed the relaxation filter; the remaining trajectories were rejected due to low AF2 confidence (n = 269) or steric clashes (n = 22).

**RFpeptides** [24] uses denoising diffusion for macrocyclic peptide binder design. Eight candidates of 14 residues were generated targeting the BMPR1A ECD. In total, 192 candidate peptides were designed across the four tools.

### Control peptides

Two types of negative controls were generated with a fixed random seed (42) for reproducibility. *Scrambled decoys* (n = 48) were produced by randomly shuffling the amino acid sequences of designed candidates (20 from PepMLM, 20 from RFdiffusion, 8 from RFpeptides), preserving amino acid composition but disrupting any sequence-dependent binding potential. BindCraft candidates were excluded from scrambling because they were generated in a separate round after control generation. *Random baselines* (n = 50) were generated with uniform amino acid sampling at lengths drawn from the empirical distribution of candidate sequences (14–25 residues). Together with the 192 designed candidates, the complete dataset comprised 290 peptides.

### Structure prediction and scoring

All 290 peptide-BMPR1A complexes were submitted to the AlphaFold 3 Server [27] for structure prediction. Jobs were distributed across five accounts (30 jobs per day per account) over two submission rounds: 266 peptides (PepMLM, RFdiffusion, RFpeptides, and all controls) in round 1, and 24 BindCraft designs in round 2. Model seeds were not specified (empty seed array), allowing the server to select random seeds and generate five independent models per job; the model with the highest ranking score was selected for downstream analysis. Extracted metrics included the interface predicted TM-score (ipTM) [32], overall ranking score, mean interface predicted aligned error (PAE_interface_), mean predicted local distance difference test (pLDDT) for the peptide chain, and the number of confident inter-chain contacts (contact probability *>* 0.5).

Orthogonal physics-based energy scoring was performed with two independent methods. **PyRosetta** [33] was used with the REF2015 energy function [34]. Each AlphaFold 3-predicted complex was subjected to constrained FastRelax minimization (200 iterations; coordinate constraint standard deviation 0.5 Å; backbone and side-chain degrees of freedom enabled). The InterfaceAnalyzerMover then computed the binding free energy (*dG*_separated_ in Rosetta energy units, REU), buried surface area (ΔSASA), interface packing statistics, and the number of unsatisfied hydrogen bonds. **FoldX** (v5.1, build 20270131) [35] was applied after converting AlphaFold 3 CIF outputs to PDB format via BioPython. Structures were first optimized with the RepairPDB command (rotamer optimization and clash repair), then scored with the AnalyseComplex command to obtain the binding free energy (Δ*G* in kcal/mol), decomposed into van der Waals, side-chain hydrogen bond, and electrostatic contributions. Energy scoring was completed for all 290 complexes with both methods.

### Contact recapitulation analysis

To assess whether designed peptides target the native BMP-2 binding site on BMPR1A, we computed contact recapitulation against the gold-standard interface from 1REW [28]. For each AlphaFold 3-predicted complex, BMPR1A residues within 5.0 Å of any peptide atom were identified using BioPython’s NeighborSearch (standard amino acids only). The recapitulation fraction was defined as the number of predicted contacts overlapping with the 30 gold-standard BMPR1A interface residues divided by 30. Novel contacts (predicted but not in the gold standard) were also recorded.

### Physicochemical property analysis

Sequence-based physicochemical properties were computed for all 290 peptides: grand average of hydropathy (GRAVY) using the Kyte-Doolittle scale, net charge at pH 7.4 via the Henderson-Hasselbalch equation with standard p*K*_*a*_ values, molecular weight (sum of residue masses minus 18.015 Da per peptide bond), isoelectric point (bisection method, 100 iterations), and the Guruprasad instability index using the DIWV dipeptide weight matrix. Candidates with | net charge | *>* 8, GRAVY *>* 0.5, or instability index *>* 40 were excluded from the filtered shortlist but retained in the full dataset for cross-tool statistical comparisons.

### Statistical analysis

Non-parametric tests were used throughout due to non-normal score distributions. The Kruskal-Wallis H-test was applied across six groups (four design tools plus two control types) for each scoring metric. Pairwise post-hoc comparisons used the Mann-Whitney U test with Bonferroni correction. Effect size *r* was computed as *r* = 1 − 2*U/*(*n*_1_ × *n*_2_). Designed candidates (n = 192) were compared against pooled controls (n = 98) using the Mann-Whitney U test. Spearman rank correlations were computed between all scoring metrics. A significance threshold of *α* = 0.05 was used throughout. All statistical analyses were performed using SciPy.

Composite ranking was computed as the mean of four per-metric ranks: ipTM (descending), PyRosetta *dG*_separated_ (ascending, more negative = stronger binding), FoldX Δ*G* (ascending), and contact recapitulation (descending, higher fraction = better interface targeting).

## Results

An overview of the computational pipeline is shown in Fig 1.

**Fig 1.**
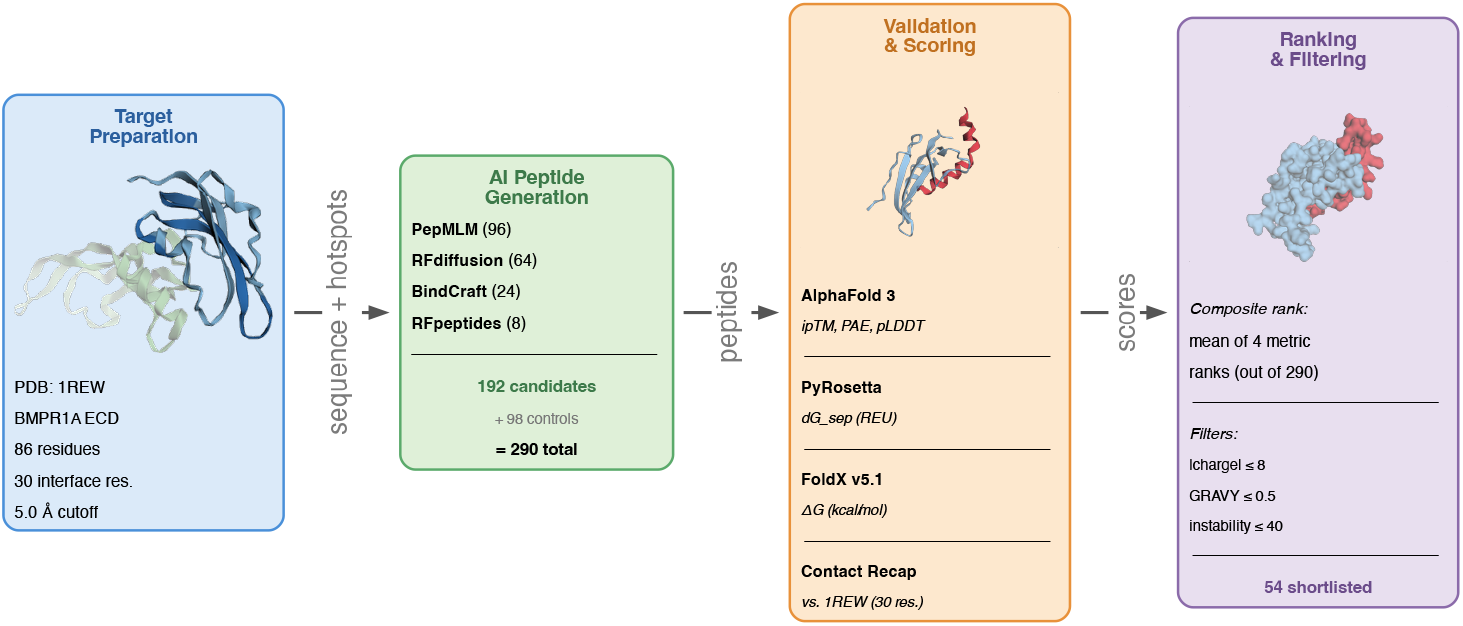
Computational pipeline overview. The study followed five stages: (1) target preparation from the 1REW crystal structure of the BMP-2:BMPR1A complex, identifying 30 interface residues at 5.0 Å cutoff; (2) AI peptide generation using four tools with distinct architectures, producing 192 candidates plus 98 controls (290 total); (3) structure prediction and scoring via AlphaFold 3, PyRosetta, FoldX, and contact recapitulation against the native interface; (4) composite ranking using all four metrics followed by physicochemical filtering, yielding 54 shortlisted candidates. Structure thumbnails show the 1REW target complex (left), an AlphaFold 3-predicted peptide–BMPR1A complex (center), and a surface representation of the top-ranked candidate (right). BMPR1A is shown in blue; peptides in red.

### Target interface characterization

The BMPR1A extracellular domain extracted from 1REW comprised 86 residues. Contact analysis at a 5.0 Å cutoff identified 30 BMPR1A residues and 20 BMP-2 residues at the binary interface, consistent with the “wrist” epitope binding geometry previously described for the BMP-2:BMPR1A interaction [28]. These 30 receptor-side interface residues served as the gold-standard contact set for evaluating whether designed peptides target the native binding site.

### Candidate generation summary

The four AI tools produced 192 candidate peptides of varying lengths and architectures (Table 1). PepMLM contributed the largest pool (n = 96; 15, 20, and 25 residues), followed by RFdiffusion (n = 64; 19 residues), BindCraft (n = 24; 11–30 residues), and RFpeptides (n = 8; 14 residues). Together with 48 scrambled and 50 random controls, the complete dataset comprised 290 peptide–BMPR1A complexes submitted to AlphaFold 3 for structure prediction and scoring.

### AlphaFold 3 validation

AlphaFold 3 structure prediction metrics varied substantially across tools (Table 2; Fig 2). BindCraft achieved the highest mean ipTM (0.595 ± 0.157) and contained the single best-scoring candidate in the study (ipTM = 0.81). PepMLM (0.547 ± 0.092) and RFdiffusion (0.546 ± 0.117) performed comparably, while RFpeptides yielded the lowest mean ipTM (0.450 ± 0.060). The same tool-level ordering was reflected in the overall ranking score and mean interface PAE, with BindCraft showing the highest confidence (7.62 ± 2.71) and RFpeptides the lowest (11.35 ± 1.38). BindCraft candidates also exhibited the highest mean peptide pLDDT (73.5 ± 9.3), followed by RFdiffusion (71.6 ± 5.5), PepMLM (59.5 ± 6.4), and RFpeptides (57.4 ± 4.7), suggesting that structure-aware design methods produce peptide conformations that AlphaFold 3 predicts with greater confidence.

**Table 2.** AlphaFold 3 structure prediction metrics by source tool. All 290 peptide–BMPR1A complexes were scored; the best model per job (by ranking score) was selected. Values are mean ± SD. Higher ipTM and ranking score indicate stronger predicted binding; lower PAE indicates higher confidence. pLDDT reflects predicted local confidence of the peptide chain.

| Tool | n | ipTM | ipTM max | Ranking score | PAE interface | pLDDT (peptide) |
| --- | --- | --- | --- | --- | --- | --- |
| BindCraft | 24 | 0.595 ± 0.157 | 0.81 | 0.694 ± 0.146 | 7.62 ± 2.71 | 73.5 ± 9.3 |
| RFdiffusion | 64 | 0.546 ± 0.117 | 0.79 | 0.682 ± 0.108 | 8.17 ± 2.04 | 71.6 ± 5.5 |
| PepMLM | 96 | 0.547 ± 0.092 | 0.74 | 0.686 ± 0.090 | 9.13 ± 1.85 | 59.5 ± 6.4 |
| RFpeptides | 8 | 0.450 ± 0.060 | 0.51 | 0.578 ± 0.052 | 11.35 ± 1.38 | 57.4 ± 4.7 |
| Scrambled ctrl | 48 | 0.515 ± 0.120 | 0.76 | 0.654 ± 0.109 | 9.92 ± 2.67 | 60.5 ± 8.1 |
| Random ctrl | 50 | 0.495 ± 0.119 | 0.83 | 0.639 ± 0.107 | 10.54 ± 2.42 | 55.6 ± 8.9 |
| **All candidates** | **192** | **0.549 ± 0.112** |  | **0.681 ± 0.105** |  |  |
| **All controls** | **98** | **0.505 ± 0.120** |  | **0.646 ± 0.108** |  |  |

**Fig 2.**
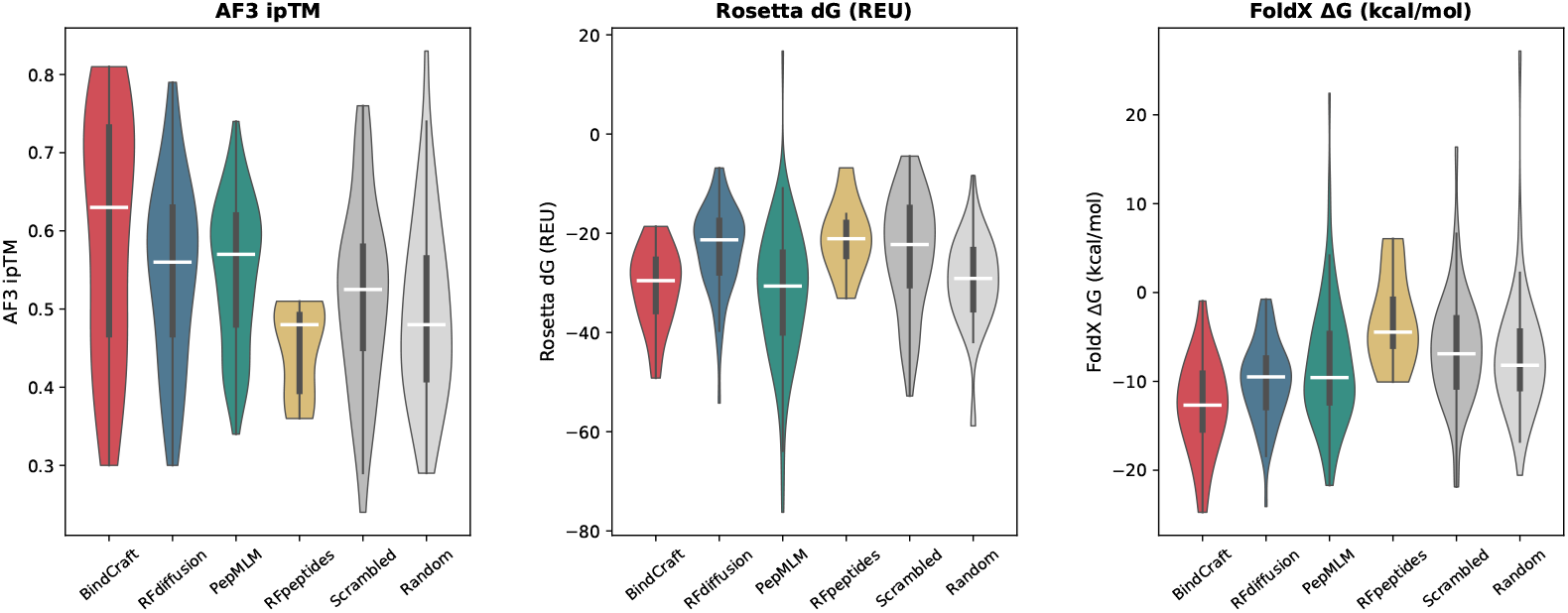
Distribution of AlphaFold 3 and energy scoring metrics by design tool. Violin plots of (A) interface predicted TM-score (ipTM), (B) PyRosetta binding free energy (dG_separated_, REU), and (C) FoldX binding free energy (ΔG, kcal/mol) for 290 peptide–BMPR1A complexes grouped by source tool. White lines indicate medians. BindCraft achieved the highest median ipTM; PepMLM the most favorable median PyRosetta dG; BindCraft and RFdiffusion the strongest FoldX ΔG. More negative energy values indicate stronger predicted binding.

Pooled designed candidates (n = 192) achieved a significantly higher mean ipTM (0.549 *±* 0.112) than pooled controls (n = 98; 0.505 *±* 0.120; Mann–Whitney *U* = 11,538, *p* = 0.002, | *r* | = 0.23). However, one random control achieved the highest individual ipTM in the entire dataset (0.83), indicating that stochastic sequences can occasionally produce high-confidence predictions.

### Energy scoring

Physics-based binding energy estimates provided orthogonal validation of AlphaFold 3 predictions (Table 3). By PyRosetta *dG*_separated_, PepMLM produced the most favorable mean binding energy (− 32.2 ± 13.6 REU), followed by BindCraft (− 30.7 ± 8.1 REU). By FoldX Δ*G*, BindCraft was strongest (− 12.8 ± 5.4 kcal/mol), followed by RFdiffusion (− 10.0 ± 4.5 kcal/mol). RFpeptides ranked last on both metrics (PyRosetta: − 20.8 ± 7.7 REU; FoldX: − 3.1 ± 5.5 kcal/mol). Notably, random controls (− 29.8 ± 9.4 REU) outperformed scrambled controls (− 24.1 ± 11.7 REU) by PyRosetta, suggesting that amino acid composition alone does not determine computed binding energy.

**Table 3.**
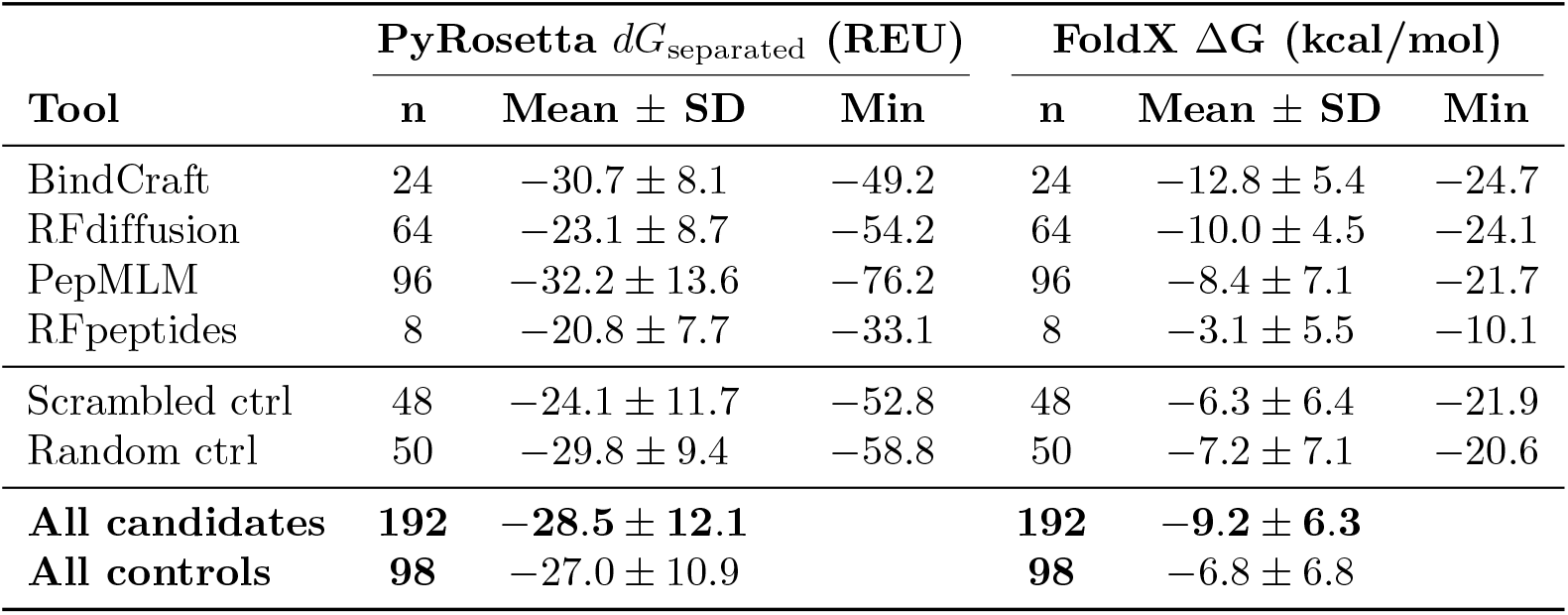
Physics-based energy scores by source tool. PyRosetta: constrained FastRelax followed by InterfaceAnalyzer (REF2015 score function; values in Rosetta Energy Units, REU). FoldX: RepairPDB relaxation followed by AnalyseComplex (values in kcal/mol). Values are mean ± SD; Min indicates the most favorable (most negative) score per tool. More negative values indicate stronger predicted binding.

Spearman correlation analysis (Fig 3) revealed a strong negative correlation between ipTM and PAE_interface_ (*ρ* = − 0.915, *p <* 10^−100^), confirming internal consistency of AlphaFold 3 outputs. The two physics-based methods showed moderate agreement (*ρ* = − 0.505, *p <* 10^−19^). Crucially, ipTM correlated only weakly with PyRosetta *dG*_separated_ (*ρ* = − 0.205) and FoldX Δ*G* (*ρ* = −0.231), indicating that the deep learning and physics-based approaches capture partially independent aspects of binding quality.

**Fig 3.**
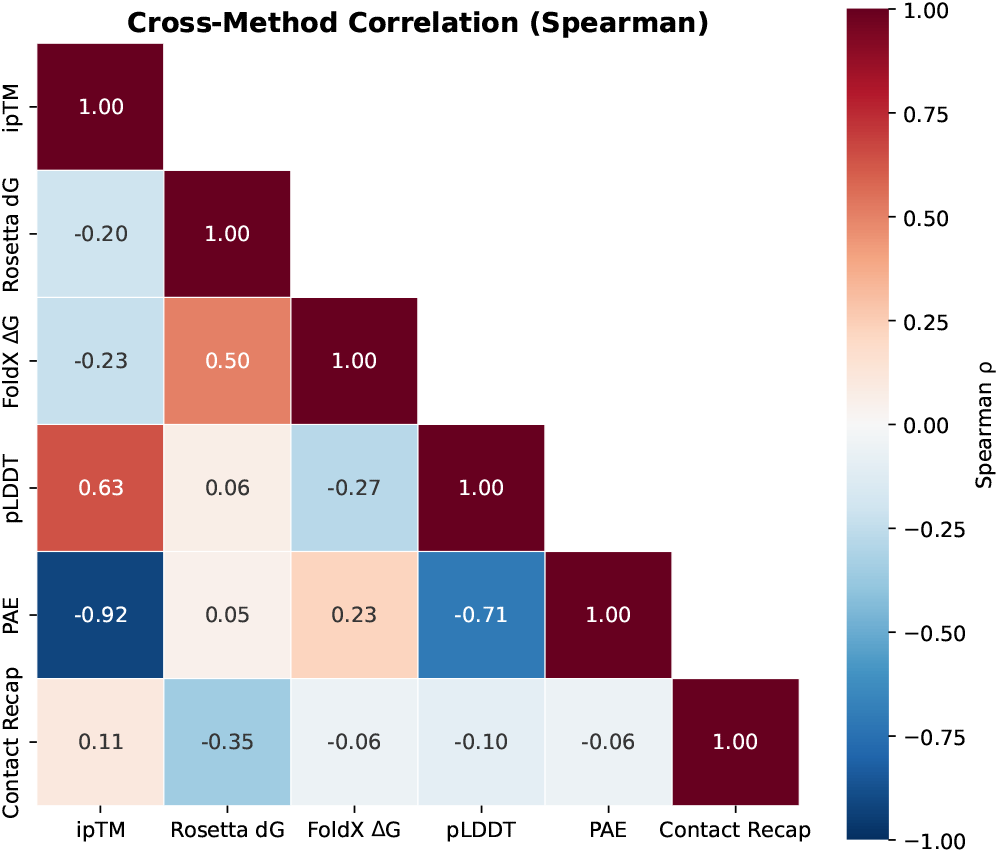
Cross-method Spearman rank correlation matrix. Pairwise Spearman correlations (*ρ*) among six scoring metrics for all 290 peptides. ipTM and PAE_interface_ are strongly anti-correlated (*ρ* = − 0.915), confirming AlphaFold 3 internal consistency. PyRosetta and FoldX show moderate agreement (*ρ* = 0.505). ipTM correlates weakly with both physics-based energies (| *ρ* | *<* 0.25), supporting their use as independent validation metrics. Contact recapitulation correlates moderately with PyRosetta dG (*ρ* = −0.352) but weakly with ipTM and FoldX.

### Contact recapitulation

Contact recapitulation was computed for all 290 peptides against the 1REW gold standard (Fig 4). PepMLM achieved the highest mean recapitulation fraction (0.249 ± 0.073), with its best candidate contacting 14 of 30 gold-standard interface residues (fraction = 0.467). RFpeptides (0.233 ± 0.036), random controls (0.227 ± 0.086), and BindCraft (0.224 ± 0.075) performed comparably, followed by RFdiffusion (0.194 ± 0.068) and scrambled controls (0.194 ± 0.070). BindCraft’s best candidate contacted 12 of 30 interface residues (fraction = 0.400). Between-group differences were significant (Kruskal–Wallis *H* = 24.91, *p* = 1.5 × 10^−4^; Table 4), but the overall comparison of designed candidates versus controls was not (Mann–Whitney *p* = 0.093). These results are consistent with the hypothesis that PepMLM—the only sequence-only tool—may preferentially sample the native interface, while structure-aware methods including BindCraft and RFdiffusion may explore alternative binding modes on the BMPR1A surface; however, the non-significant pooled comparison (*p* = 0.093) warrants caution in this interpretation.

**Table 4.**
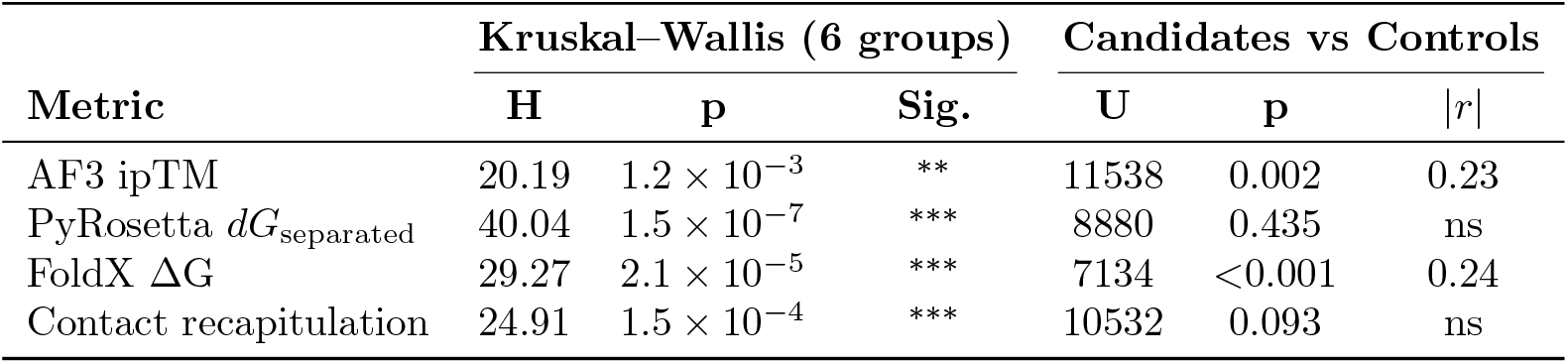
Statistical comparisons across design tools and between candidates and controls. Kruskal–Wallis H-tests assessed differences across all six groups (four tools plus two control types); Mann–Whitney U tests compared pooled designed candidates (n = 192) versus pooled controls (n = 98). Effect size | *r* | is computed as | 1 − 2*U/*(*n*_1_*n*_2_) |; values ≥ 0.20 indicate a small-to-medium effect. ^∗^*p <* 0.05; ^∗∗^*p <* 0.01; ^∗∗∗^*p <* 0.001; ns, not significant.

**Fig 4.**
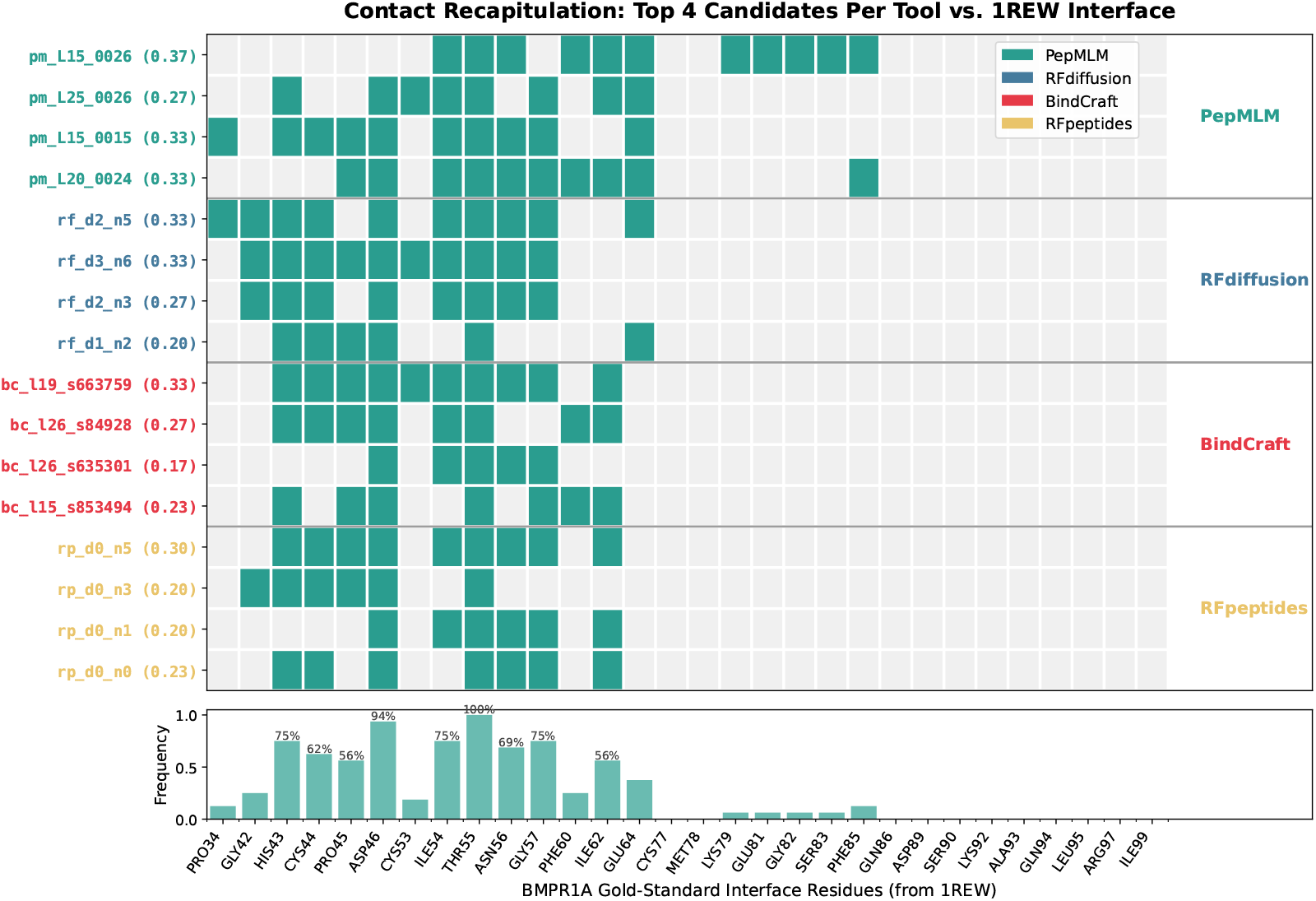
Contact recapitulation heatmap of top candidates against the 1REW gold-standard interface. Binary heatmap showing which of the 30 BMPR1A gold-standard interface residues (columns) are contacted by the top 4 designed candidates from each tool (rows), at a 5.0 Å cutoff. Contact fractions are shown in parentheses. Horizontal lines separate tool groups. The bottom panel shows the frequency with which each residue is contacted across all 16 candidates. Core hotspot residues Asn55–Gly57 are contacted by nearly all top candidates (≥ 87%), while PepMLM candidates show broader coverage extending to C-terminal interface residues (Glu64–Gln86).

### Cross-tool statistical comparison

The Kruskal–Wallis test detected significant between-group heterogeneity for all four primary metrics (Table 4): ipTM (*H* = 20.19, *p* = 1.2 × 10^−3^), PyRosetta *dG*_separated_ (*H* = 40.04, *p* = 1.5 × 10^−7^), FoldX Δ*G* (*H* = 29.27, *p* = 2.1 × 10^−5^), and contact recapitulation (*H* = 24.91, *p* = 1.5 × 10^−4^).

When designed candidates were compared against controls as two groups (Fig 5), significant advantages were observed for ipTM (*p* = 0.002, |*r*| = 0.23) and FoldX Δ*G* (*p <* 0.001, | *r* | = 0.24), but not for PyRosetta *dG*_separated_ (*p* = 0.435) or contact recapitulation (*p* = 0.093). The small-to-medium effect sizes suggest that current generative AI tools produce a measurable but modest improvement over random baselines in computational binding metrics.

**Fig 5.**
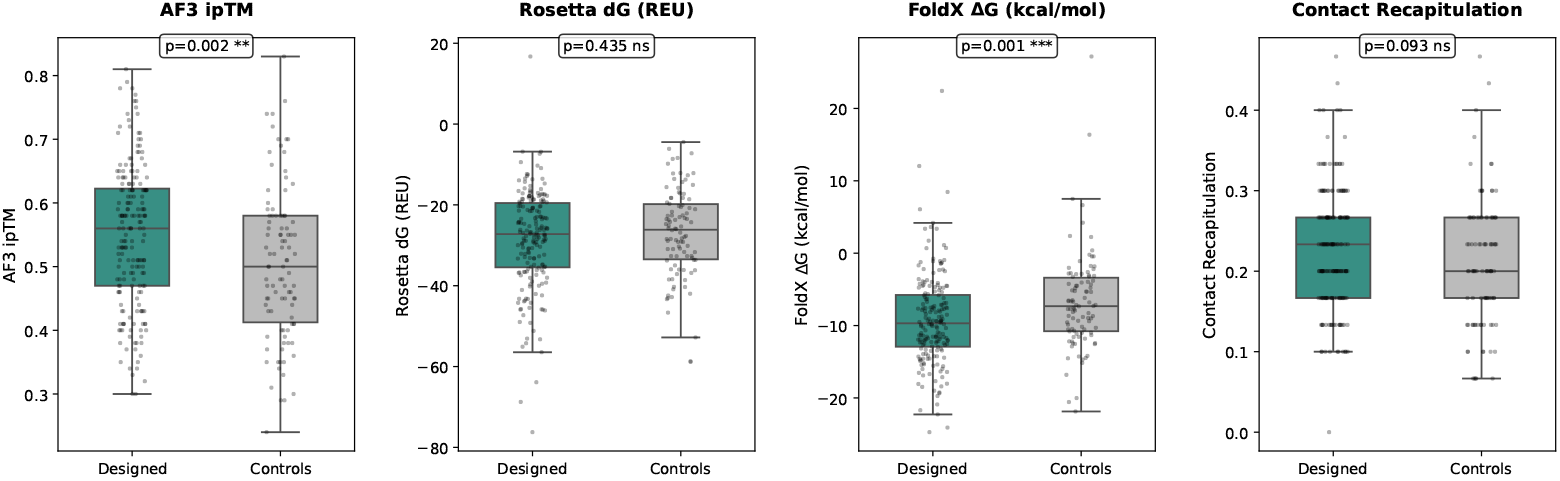
Designed candidates versus controls across four scoring metrics. Box-and-strip plots comparing 192 designed candidates against 98 controls (48 scrambled + 50 random) for (A) AF3 ipTM, (B) PyRosetta dG_separated_, (C) FoldX ΔG, and (D) contact recapitulation. *p*-values from two-sided Mann–Whitney U tests are shown. Designed candidates significantly outperformed controls on ipTM (*p* = 0.002) and FoldX ΔG (*p <* 0.001), but not on PyRosetta dG (*p* = 0.435) or contact recapitulation (*p* = 0.093).

### Top candidates

Composite ranking integrated ipTM, PyRosetta *dG*_separated_, FoldX Δ*G*, and contact recapitulation into a single score per peptide (Table 5; Fig 6). The top-ranked candidate was pepmlm L15 0026 (composite rank 19.8; ipTM = 0.66, PyRosetta *dG*_separated_ = − 45.9 REU, FoldX Δ*G* = − 19.4 kcal/mol, contact fraction = 0.367), a 15-residue PepMLM design that combined strong energy scores with the highest contact recapitulation among top candidates. The second-ranked candidate, rfdiff_d2_n5 (rank 20.5; ipTM = 0.79, contact = 0.333), achieved the highest ipTM among the top 10. When all 290 peptides were ranked, two controls appeared in the overall top 10: random_0045 (rank 33.0, position 4) and scrambled_PepMLM_0016 (rank 46.0, position 10), underscoring the need for experimental validation. Among designed candidates only (Table 5), the top 10 comprised six PepMLM, three RFdiffusion, and one BindCraft peptide, spanning lengths from 15 to 25 residues. No RFpeptides candidates appeared in the top 10. Notably, BindCraft candidates—despite achieving the highest mean ipTM (0.595)—ranked lower in the four-metric composite due to moderate contact recapitulation (mean 0.224), raising the possibility that they bind alternative sites on the BMPR1A surface rather than the native BMP-2 interface, though this interpretation remains a computational hypothesis pending experimental confirmation.

**Table 5.**
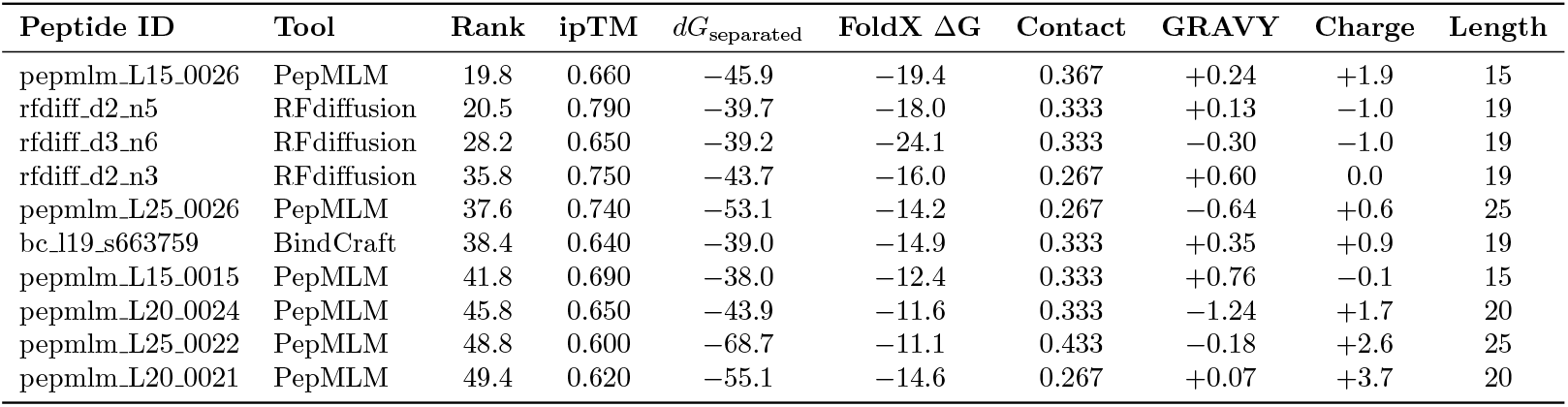
Top 10 designed peptide candidates ranked by four-metric composite score. The composite is the mean of each peptide’s individual ranks (out of 290) on AF3 ipTM, PyRosetta *dG*_separated_, FoldX ΔG, and contact recapitulation; lower values indicate consistently high performance across all four metrics. Contact: fraction of 30 gold-standard BMPR1A interface residues contacted at 5.0 Å. GRAVY: grand average of hydropathy (Kyte-Doolittle); Charge: net charge at pH 7.4. Controls are excluded from this table; two controls appeared in the overall top 10 (see text).

**Fig 6.**
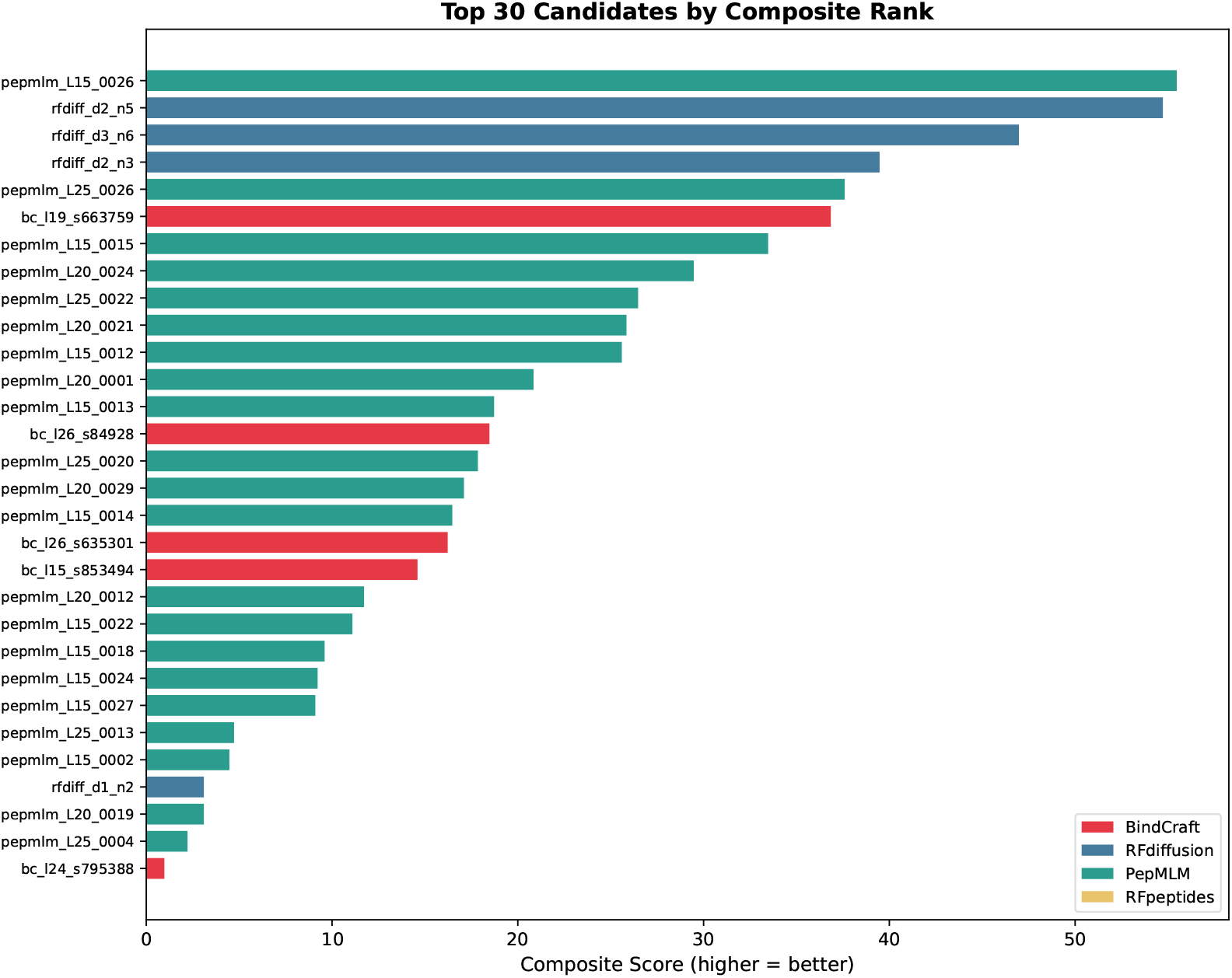
Top 30 designed candidates ranked by four-metric composite score. Horizontal bar chart showing the top 30 designed peptides (controls excluded) ranked by composite score, computed as the mean of per-peptide ranks on ipTM, PyRosetta dG_separated_, FoldX ΔG, and contact recapitulation (out of 290). Higher bars indicate better composite performance. Colors indicate the design tool of origin. PepMLM candidates dominate the top ranks, reflecting their balanced performance across binding confidence, energy, and native interface targeting.

Physicochemical filtering (Fig 7) removed candidates with extreme net charge (| charge | *>* 8), high hydrophobicity (GRAVY *>* 0.5), or predicted instability (instability index *>* 40), yielding a filtered shortlist of 54 designed candidates: PepMLM (22), RFdiffusion (16), BindCraft (9), and RFpeptides (7). Twenty-eight controls (16 random, 12 scrambled) also satisfied all three physicochemical criteria, yielding similar pass rates for designed (54/192, 28.1%) and control (28/98, 28.6%) peptides. This is expected, as physicochemical filters assess sequence composition rather than binding competence; the shortlist was restricted to designed candidates for downstream experimental prioritization. Among the top 10 designed candidates, one (rfdiff_d2_n3, GRAVY = +0.60) was excluded from the shortlist due to elevated hydrophobicity.

**Fig 7.**
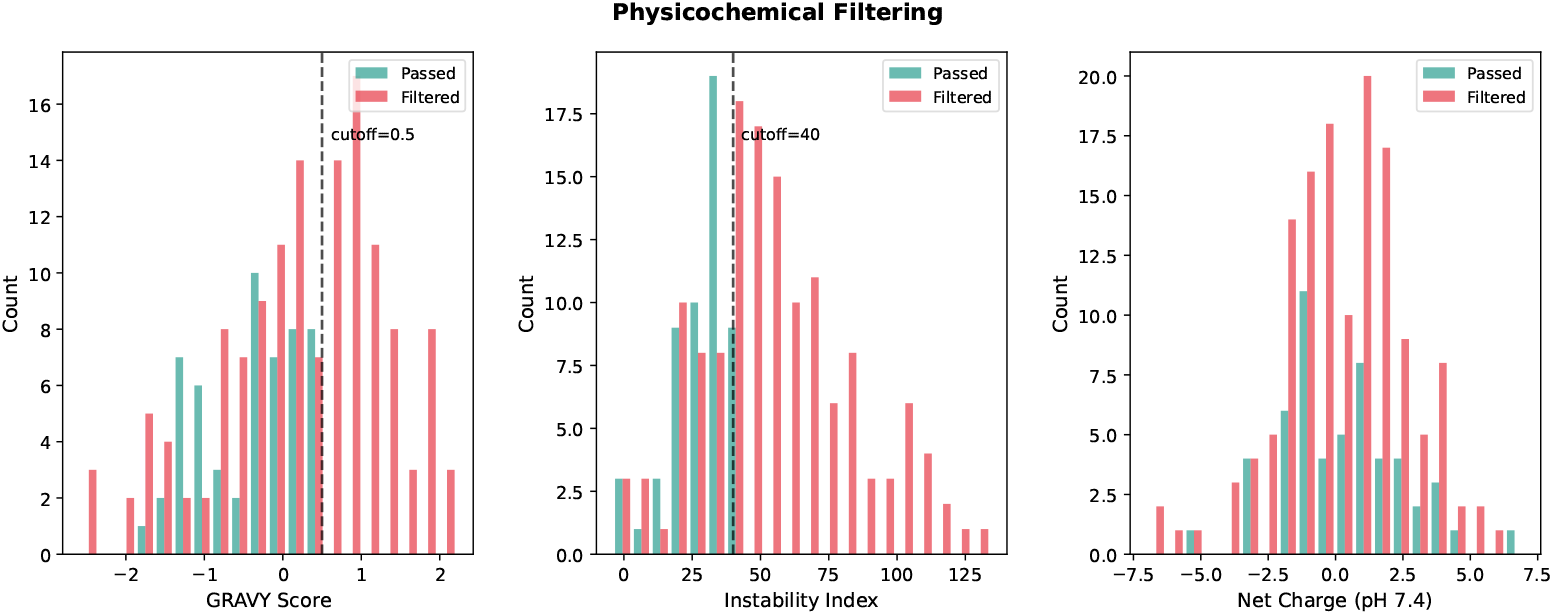
Physicochemical property distributions of designed candidates. Histograms of (A) GRAVY hydrophobicity, (B) Guruprasad instability index, and net charge at pH 7.4 for 192 designed candidates, colored by filter outcome: passed all three criteria (teal) or excluded (red). Dashed lines indicate filter cutoffs (GRAVY *>* 0.5; instability index *>* 40). Candidates with | charge | *>* 8, GRAVY *>* 0.5, or instability index *>* 40 were excluded from the filtered shortlist of 54 candidates.

Structural analysis of three representative candidates from distinct tools (Fig 8) revealed different binding geometries. The top-ranked PepMLM candidate (L15_0026, 15 residues) showed a partially helical structure with broad interface coverage (11/30 gold-standard residues, contact = 0.37). The second-ranked RFdiffusion candidate (d2_n5, 19 residues) adopted a compact single-helix conformation with the highest ipTM among the top 10 (0.79) and 10/30 interface contacts. The highest-ipTM BindCraft candidate (l26_s635301, 26 residues; ipTM = 0.81) formed an extended helix-turn-helix motif but contacted only 5/30 gold-standard residues (contact = 0.17), illustrating the dissociation between ipTM and native interface targeting that motivates the four-metric composite.

**Fig 8.**
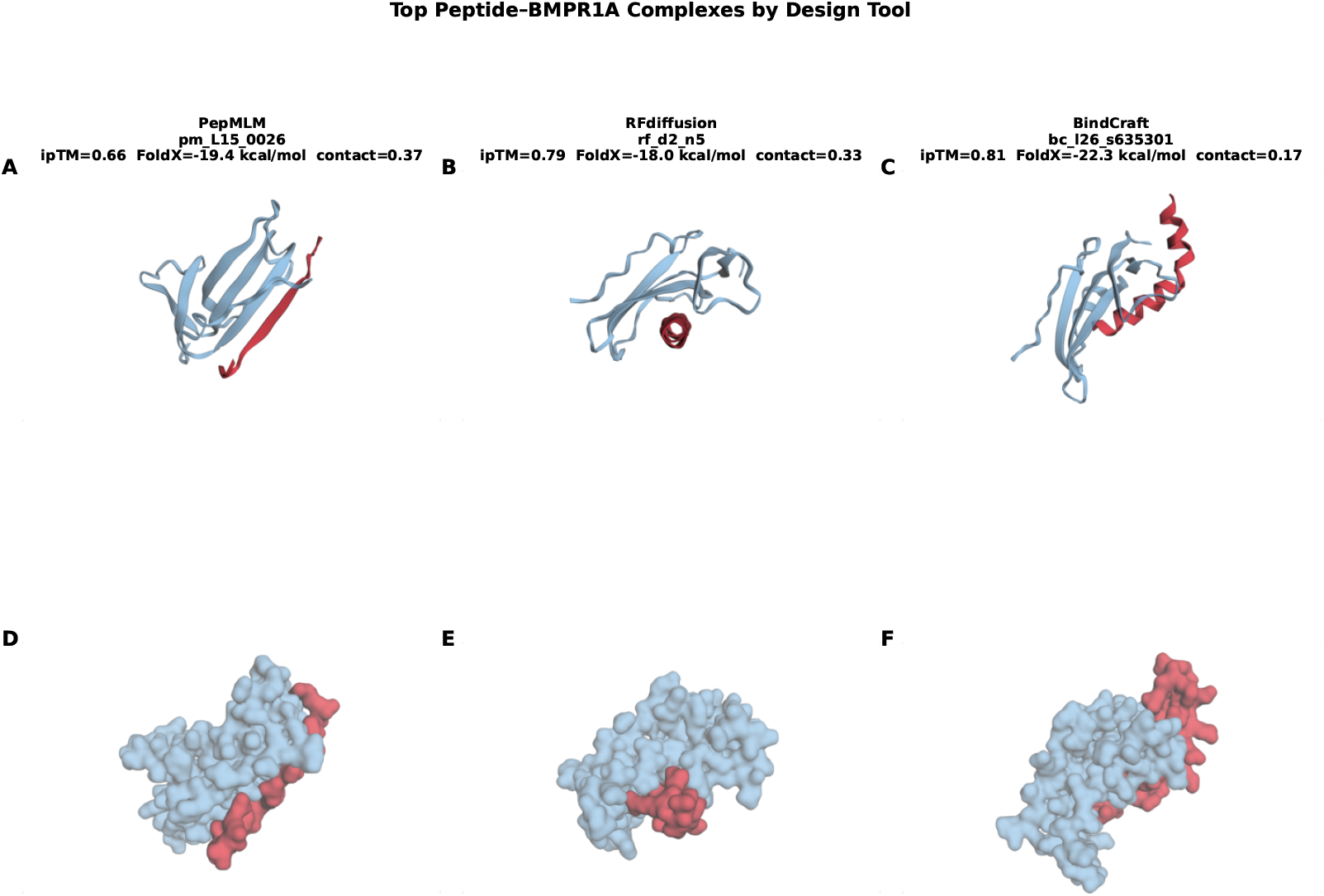
AlphaFold 3-predicted structures of representative peptide–BMPR1A complexes. Cartoon (A–C) and surface (D–F) representations of three candidates from distinct design tools. (A, D) PepMLM candidate L15_0026 (composite rank 1; ipTM = 0.66, contact = 0.37): partially helical peptide with broad interface coverage. (B, E) RFdiffusion candidate d2_n5 (rank 2; ipTM = 0.79, contact = 0.33): compact single-helix conformation. (C, F) BindCraft candidate l26_s635301 (highest ipTM = 0.81, contact = 0.17): extended helix-turn-helix motif binding an alternative BMPR1A site. BMPR1A shown in light blue; peptides in red. The BindCraft candidate illustrates the dissociation between high ipTM and low native interface targeting.

## Discussion

This study represents, to our knowledge, the first application of generative AI protein design tools to cartilage regeneration targets. By benchmarking four architecturally distinct methods—RFdiffusion, BindCraft, PepMLM, and RFpeptides—against BMPR1A using a four-metric composite ranking, we identified 54 shortlisted candidates and a top-ranked 15-residue PepMLM design (pepmlm_L15_0026) that combines favorable binding energy with the highest contact recapitulation among leading candidates. The study also revealed an unexpected dissociation between structural confidence and native interface targeting: BindCraft candidates achieved the highest mean ipTM (0.595) yet exhibited moderate contact recapitulation (0.224), raising the possibility that they engage alternative binding surfaces on the BMPR1A extracellular domain.

The four AI architectures exhibited complementary strengths that reflect their underlying design principles. PepMLM, the only sequence-conditioned method in our panel, dominated the composite ranking (six of the top 10 designed candidates) and achieved the highest mean contact recapitulation (0.249), consistent with its training on natural protein–peptide interfaces that may bias it toward evolutionarily conserved binding modes. RFdiffusion produced candidates with the highest ipTM among non-BindCraft tools (0.79 max) and the most favorable FoldX energies, reflecting the strength of diffusion-based backbone generation for producing physically plausible binding geometries. BindCraft candidates showed the highest structural confidence overall (ipTM 0.81 max, pLDDT 73.5 mean), consistent with its AlphaFold 2-guided optimization, but their lower contact recapitulation scores suggest possible engagement of non-native sites, highlighting the risk of optimizing for a confidence metric that does not enforce epitope specificity. RFpeptides underperformed across all metrics, possibly because its macrocyclic design constraints are less suited to the relatively flat BMPR1A wrist epitope, which may favor extended or helical peptide conformations over cyclic scaffolds.

Our AI-designed candidates represent a fundamentally different approach from existing BMP mimetic peptides. Natural BMP-2 knuckle epitope peptides, while capable of inducing bone formation [15, 16], exhibit 12,000-fold lower osteoinductive potency than the full-length protein and suffer from aggregation [19]. CK2.1, the BMPR1A mimetic peptide that achieved cartilage repair in an OA mouse model [9], acts intracellularly by mimicking the receptor’s intracellular domain rather than binding the extracellular domain. Our peptides are designed to bind the BMPR1A ECD at the native BMP-2 interface, potentially functioning as competitive agonists or antagonists—a mechanism that would need to be resolved experimentally. Emerging tools such as PepMimic [36], which designs peptide binders by mimicking known protein–protein interfaces, could complement our approach by leveraging the structural data generated in this study.

Our validation strategy was designed to mitigate the known limitations of any single computational metric. AlphaFold 3 ipTM served as the primary measure of binding confidence [27, 32], but benchmarking studies have shown that AlphaFold 3 complex structures can exhibit large errors not captured by ipTM, particularly for flexible regions [37]. By combining ipTM with two independent physics-based energy functions (PyRosetta and FoldX) and contact recapitulation against a crystallographic reference, we reduced the probability that a top-ranked candidate succeeds on one metric by chance. The weak correlation between ipTM and both energy metrics (| *ρ* | *<* 0.25) confirms that these approaches capture partially independent aspects of binding quality. Nevertheless, all four metrics remain computational proxies: none directly measures binding affinity, and high-scoring candidates may still fail experimentally [26].

Several limitations should be noted. First, no experimental validation was performed; all conclusions are based on computational predictions. Second, the study targets a single receptor (BMPR1A), and the relative tool performance may differ for other targets with different surface geometries. Third, our analysis cannot distinguish agonists from antagonists—a peptide that binds the BMP-2 epitope on BMPR1A could either activate or inhibit downstream signaling depending on whether it induces the conformational changes required for type II receptor recruitment. Fourth, the short peptide length constraint (11–25 residues for most tools; 14 for RFpeptides) may disadvantage tools optimized for larger protein binders. Fifth, scrambled controls retain the amino acid composition of their parent sequences, which may partially preserve physicochemical properties relevant to non-specific binding, potentially narrowing the observed differences between designed and control peptides. Sixth, AlphaFold 3 Server predictions were obtained in two submission rounds (266 peptides in round 1, 24 BindCraft designs in round 2) with unspecified random seeds; although we selected the best of five models per job to reduce stochastic variation, we cannot rule out minor server-side differences between rounds, and the BindCraft-specific findings should be interpreted with this caveat.

Several directions could advance this work toward therapeutic application. Molecular dynamics simulations of the top candidates in explicit solvent would assess binding stability and identify candidates whose interactions persist over nanosecond timescales. Experimental validation should prioritize surface plasmon resonance (SPR) or isothermal titration calorimetry (ITC) binding assays against purified BMPR1A ECD, followed by functional assays in chondrocyte micromass cultures to determine whether candidate peptides act as agonists or antagonists of BMP signaling. Extending the design to the ternary BMP-2:BMPR1A:ActRII complex could yield peptides that modulate receptor heterodimerization. Finally, the computational framework established here—multi-tool generation, multi-metric validation, and contact recapitulation against a crystallographic reference—is generalizable to other receptor targets in regenerative medicine.

## Conclusion

This study demonstrates that current generative AI tools can produce candidate BMPR1A-binding peptides with computational binding signatures that significantly exceed random baselines on multiple independent metrics. The four-metric composite ranking identified a shortlist of 54 candidates spanning four architectural paradigms, with PepMLM designs achieving the strongest balanced performance across structural confidence, binding energy, and native interface targeting. The observation that BindCraft candidates exhibit lower contact recapitulation despite high ipTM scores underscores the importance of contact recapitulation as a validation metric alongside confidence-based measures. While experimental validation remains essential, these results establish a reproducible computational framework for AI-guided peptide design in cartilage regeneration and identify specific candidates for future in vitro and in vivo testing.

## Data availability

All code, data, and figures are publicly available at https://github.com/shirtlessman/bmpr1a-peptide-design. The repository includes the complete computational pipeline, designed peptide sequences, AlphaFold 3 input files, scoring results, and interactive 3D structure viewers.

## Supporting information

**S1 Table. Complete scoring data for all 290 peptide-BMPR1A complexes**. Master scores table including AF3 metrics, PyRosetta energy scores, FoldX binding energies, contact recapitulation fractions, and physicochemical properties for all 192 designed candidates and 98 controls. Available at https://github.com/shirtlessman/bmpr1a-peptide-design/blob/main/data/results/master_scores.csv.

**S1 File. Designed peptide sequences**. FASTA file containing all 192 designed candidate sequences with tool of origin and design parameters. Available at https://github.com/shirtlessman/bmpr1a-peptide-design/tree/main/data/candidates.

**S1 Fig. Interactive 3D structure gallery**. HTML-based interactive visualization of top candidate peptide-BMPR1A complexes predicted by AlphaFold 3, rendered with Py3Dmol. Available at https://shirtlessman.github.io/bmpr1a-peptide-design/figures/structures/index.html.

## Acknowledgments

We thank the developers of the open-source tools that made this work possible: the Institute for Protein Design at the University of Washington for RFdiffusion, RFpeptides, and ProteinMPNN; Pacesa, Correia (EPFL), and Ovchinnikov (MIT) for BindCraft; the Chatterjee laboratory (University of Pennsylvania) for PepMLM; the PyRosetta and Rosetta Commons communities; and the FoldX team. We acknowledge Google DeepMind and Isomorphic Labs for providing public access to the AlphaFold 3 Server and Google Colab for GPU computing resources. Structure data were obtained from the RCSB Protein Data Bank. We are grateful to the broader open-source scientific computing community, including the developers of BioPython, SciPy, NumPy, pandas, matplotlib, and seaborn, whose tools underpin the analyses presented here.

